# A chemical biology toolbox to investigate in-cell target engagement and specificity of PRMT5-inhibitors

**DOI:** 10.1101/2022.01.20.477145

**Authors:** Elisabeth M. Rothweiler, Jakub Stefaniak, Jennifer A. Ward, Catherine Rogers, Esra Balikci, Kilian V. M. Huber

## Abstract

Increasing evidence suggests the protein arginine methyltransferase PRMT5 as a contributor to tumorigenesis in various cancer types and several inhibitors have entered clinical trials. Robust assays to determine cellular target engagement and selectivity are an important asset for the optimisation of inhibitors and the design of relevant *in vivo* studies. Here we report a suite of chemical biology assays enabling quantitative assessment of PRMT5 inhibitor in-cell target engagement and global selectivity profiling using a representative set of inhibitors. With the help of a bespoke cellular probe, we assess inhibitor target occupancy in cells in relation to biochemical and functional cellular assays. Investigating the influence of SAM, the natural cofactor of PRMT5, our results support the hypothesis that SAM positively contributes to the engagement of substrate-competitive inhibitors via a PRMT5:SAM:inhibitor ternary complex. Extensive proteomic profiling studies by drug affinity chromatography and thermal profiling further indicate high specificity of the clinical PRMT5 inhibitor GSK3326595 (pemrametostat).

**Graphical abstract:** 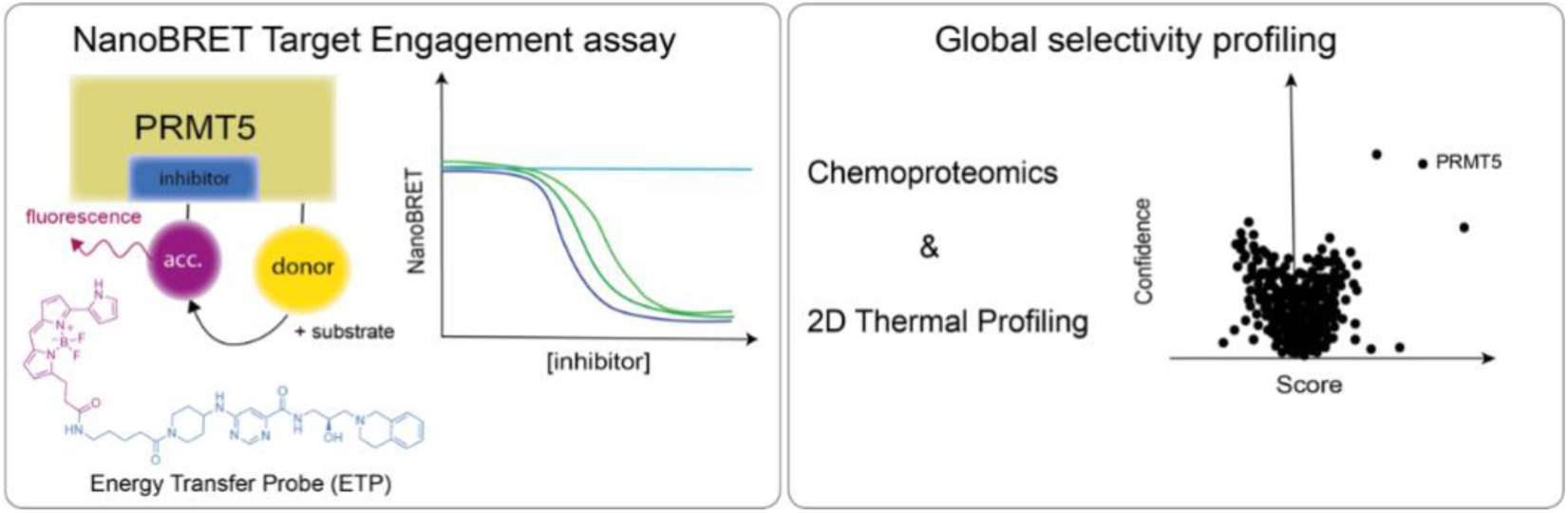

## Introduction

Protein arginine methyl transferases (PRMTs) transfer a methyl group from their cofactor S-adenosylmethionine (SAM) to protein substrates. Cofactor and substrate bind at distinct sites that are connected through a central channel where the methyl transfer takes place. The reaction results in methylarginine residues on histone and non-histone proteins and release of S-adenosyl-homocysteine (SAH). The methylation of arginine’s guanidinium group does not alter its positive charge at physiological pH but affects its hydrophobicity and ability to mediate inter- and intra-molecular interactions.^*1, 2*^ There are nine human PRMTs that are classified by their mode of methyl group transfer into PRMT family types I, II, III and IV. Type II PRMTs, PRMT5 and PRMT9, process symmetrical dimethylarginine (sDMA) by adding a methyl group to the neighbour nitrogen atom of guanidine.^*3*^ PRMT5 regulates transcription by methylating histones, transcription factors, nucleosome remodelling- and co-repressor complexes. Furthermore, it is thought to play a role in splicing regulation as part of the methylosome.^*4*^ PRMT5 distinguishes its methylation targets by interacting with nuclear and cytosolic adaptor proteins such as CLNS1A, WDR77 (MEP50), RIOK1 or COPR5.^*5*^ PRMT5 overexpression has been observed in a range of aggressive cancer types such as lymphoma, melanoma, glioblastoma, gastric and breast cancer.^*6*^ In the latter, PRMT5 has been designated as a marker for poor prognosis. PRMT5 may promote cell transformation and proliferation,^*7, 8*^ and genetic knockdown studies revealed mitigating effects on cancer cell proliferation.^*9*^ The collected evidence for its role as a promotor of growth, proliferation, migration and invasion in cancer suggests PRMT5 as an emerging target for drug discovery with both the substrate and SAM-binding sites being amenable to targeting with small molecule inhibitors (Fig. 1A).^*10, 11*^ By its nature, the substrate binding site is structurally diverse as PRMTs methylate a range of distinct proteins which in turn offers opportunities for targeted and PRMT-subtype selective inhibitor development. Pharmacological modulation via the cofactor binding pocket is more challenging as interactions with SAM are more conserved and SAM-competitive inhibitors need to be polar to access the pocket yet still hydrophobic enough to cross cellular membranes.^*12*^ However, inhibitors for both sites have been identified successfully. The development of SAM-competitive PRMT5 inhibitors such as LLY-283 was inspired by methylthioadenosine, an endogenous ligand inhibiting PRMT5 selectively.^*13*^ It is a potent nanomolar range inhibitor in biochemical assays and showed sub-micromolar potency in cells.^*14*^ GSK3326595 (pemrametostat), a highly potent substrate-competitive PRMT5 inhibitor, is currently being investigated in clinical trials.^*15*^

**Figure 1:**
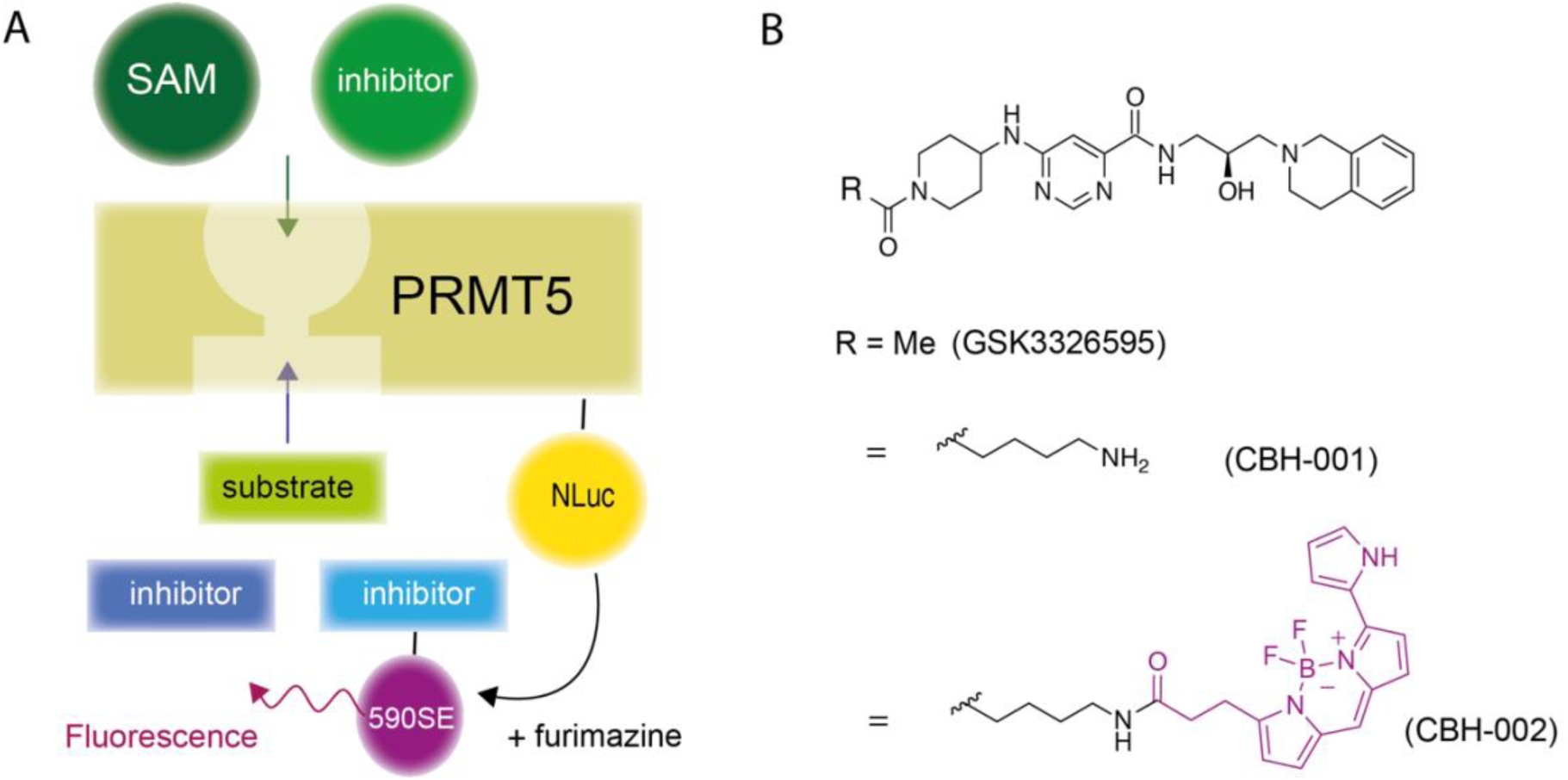
PRMT5 NanoBRET assay and ETP probe development. A) NanoBRET TE assay concept cartoon showing PRMT5 binding pockets for SAM and substrates. Upon addition of furimazine, the PRMT5-NLuc fusion protein excites the ETP’s fluorophore resulting in BRET. B) Structures of GSK3326595 and analogues CBH-001 and CBH-002.

Confirmation and quantification of in-cell target engagement (TE) is a key parameter in drug development complementing the usually acellular high-throughput hit identification and early lead optimisation stages.^*16–18*^ The increased complexity in cells, in comparison to biochemical assays using purified protein, aids in understanding more broadly the effect of pharmacologic modulation of the target of interest. This also includes the influence of posttranslational modifications, varying metabolite levels, and protein-protein interactions, all of which can be critical for biomarker identification and patient stratification.^*19*^ Examples for well-established techniques to study TE include e.g. chemoproteomics,^*20*^ cellular thermal shift assay (CETSA) ^*21*^ and bioluminescence resonance energy transfer (BRET). BRET provides a robust approach to quantify the formation of compound-target complexes in live cells. When BRET partners come into close proximity (< 10 nm), energy is transferred from a donor molecule to an acceptor moiety by Förster transfer. Small molecules can act as acceptors when conjugated to a fluorophore (often referred to as energy transfer probes (ETP)) and BRET is generated upon ETP binding to a protein of interest (POI) fused with NanoLuc (NLuc) luciferase (Fig. 1A).^*22, 23*^

Here, we present a cell permeable ETP enabling quantification of PRMT5 inhibitor TE in living cells via NanoBRET. In addition, we profile the clinical PRMT5 inhibitor GSK3326595 using complementary chemical proteomic as well as thermal profiling approaches suggesting exquisite selectivity of this substrate-competitive inhibitor in cells.

## Results

### Development of a PRMT5 energy transfer probe and NanoBRET assay optimisation

To establish a bespoke PRMT5 NanoBRET TE assay we developed a fluorescent ETP, CBH-002, based on the known substrate-competitive inhibitor GSK3326595 (Fig. 1B). Introduction of a short aminobutyl linker enabled convenient conjugation of the BRET acceptor fluorophore to the intermediate CBH-001 via activated NHS-ester amide coupling (SI).

Next, we compared N-versus C-terminal fusions of PRMT5 with NLuc to examine any potentially preferred orientation for the generation of BRET signal. Results suggested the N-tagged NLuc-PRMT5 was favoured over the C-terminal fusion (Supp. Fig. 1B). We then titrated CBH-002 against fixed concentrations of DMSO and the parent compound GSK3326595 which indicated a suitable assay window between 0.5-0.1 μM ETP (Supp. Fig. 1C). We estimated the apparent K_d_ by testing eight fixed ETP concentrations against a dilution series of GSK3326595 (Fig. 2A).^*22*^ The resulting EC_50_ values were plotted against the ETP concentrations (Fig. 2B) and linear regression analysis in combination with the Cheng-Prusoff equation^*24*^ suggested an apparent K_d_ value of 13.5 ± 6.2 nM. Ideally, the ETP concentration should provide a good assay window while resulting in stable EC_50_ values. To achieve accurate quantitation of binding, excess quantities of ETP should be avoided, as they may lead to oversaturation and ablated inhibitor binding. Conversely, suboptimal ETP concentrations may risk competition by the inhibitor. We therefore selected a concentration of 0.075 μM ETP to evaluate PRMT5 inhibitor binding as this concentration delivered stable EC_50_ values whilst maintaining an adequate assay window (Fig. 2A, Supp. Fig. 1C).

**Figure 2:**
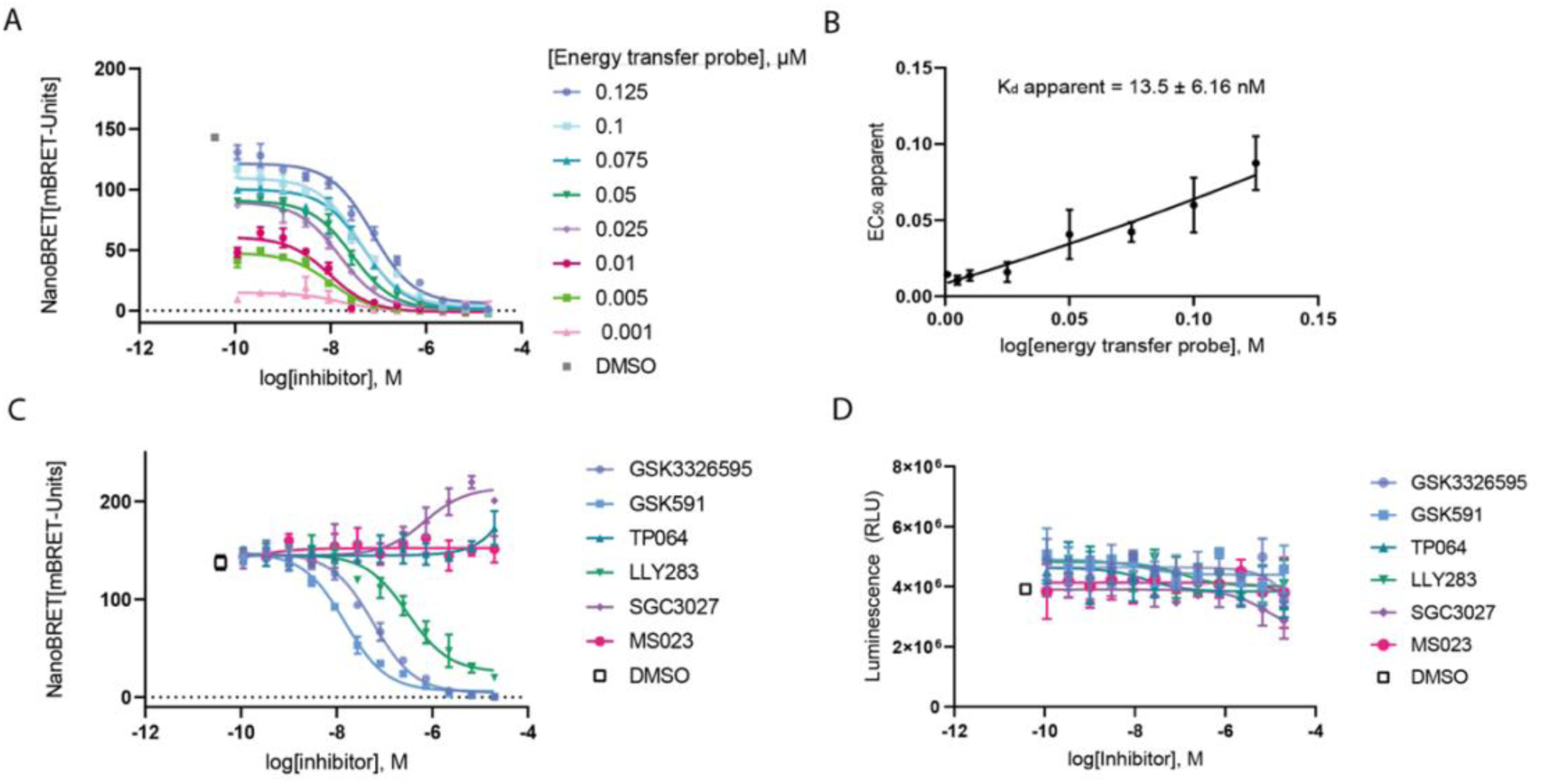
Optimisation of ETP concentration and evaluation of PRMT5 inhibitors in a cell-based NanoBRET assay using CBH-002. A) ETP titration demonstrating cellular TE (n=4). B) The apparent K_d_ was determined as the Y-intercept of compound EC_50_ values plotted against ETP concentration by linear regression. C) Comparison of inhibitor EC_50_ values in HEK293 cells transfected with NL-PRMT5 after treatment for 2 h with compound and ETP. Each data point represents the mean ± SD of three biological experiments (n=3). D) Cell viability control via CellTiterGlo assay.

### Measurement of EC_50_ values in cells

Having established optimal ETP conditions we set out to assess live cell TE of various well-known cell active PRMT inhibitors (Fig. 1B and Supp. Fig. 1A). To test the specificity of our assay, we also included inhibitors of other PRMTs such as SGC3027 which is a selective chemical probe for PRMT7.^*25*^ We further selected MS023, a potent substrate-mimetic PRMT type I inhibitor^*26*^ as well as TP064 which is a selective inhibitor for PRMT4 binding to the substrate binding site.^*27*^ When tested in our assay, TP064 and MS023 did not exhibit any significant displacement of our PRMT5 ETP, thus confirming selectivity for their respective targets (Fig. 2C). However, we noticed an increase in BRET signal for SGC3027 which appeared to correlate with a concomitant loss of signal in cell viability assays (Fig. 2D). We hypothesise this may be due to pre-apoptotic stress induced by the inhibitor. The known PRMT5 inhibitors LLY-283, GSK3203591 and GSK3326595 afforded dose-response curves in line with expectations and without any apparent effects on cell viability. Interestingly, although derived from a substrate-competitive inhibitor scaffold, CBH-002 was able to determine EC_50_ values for both substrate-as well as cofactor-competitive PRMT5 inhibitors. In comparison to acellular biochemical assays monitoring inhibition of methylation using the purified PRMT5:MEP50 complex, the intact cell NanoBRET EC_50_ values were approximately 5-to 10-fold higher for the substrate-competitive compounds GSK3326595 and GSK3203591, whereas for the SAM-competitive LLY-283 potency was markedly reduced (Table 1). Results from previously published functional cellular assays evaluating symmetric arginine methylation in various cell lines via functional in-cell Western (ICW) and ELISA indicated EC_50_ values in the range of 0.3–56 nM for GSK3203591 and GSK3326595.^*28–30*^ In case of LLY-283, an IC_50_ value of 25 nM has been reported from an assay investigating SmBB’ methylation in MCF7 cells.^*31*^ In this context, the NanoBRET EC_50_ values obtained for GSK3203591 in our HEK293 system align well with the published data. In comparison, the effective half-maximal concentrations for target engagement for GSK3326595 and LLY-283 appear higher versus these functional readouts. As displacement of the ETP by the inhibitors is indicative of binding affinity, we therefore sought to compare our results with published biophysical data. Unfortunately we could not retrieve any such data for GSK3326595 and GSK3203591, yet for the closely related compound GSK3235025 (EPZ015666) SPR K_d_ values of 171 nM and less than 1 nM have been reported for SAH- and SAM-bound PRMT5:MEP50, respectively.^*15*^ For LLY-283, a K_d_ value of approximately 6 nM has been determined for PRMT5.^*14*^ Naturally, the impact of increased complexity including any uptake or export mechanisms, and in particular varying cofactor levels across different cell lines, as well as the use of different readouts and techniques is difficult to gauge. Thus, we decided to further investigate how modulation of SAM levels might affect the different types of inhibitors in our experimental setup.

**Table 1:** Comparison of literature data and NanoBRET TE assay results. ^a^ HTRF assay monitoring monomethylation of H4R3 on H4 peptide by PRMT5:MEP50; ^b^ Radioactive assay monitoring methyl transfer from ^3^H-SAM to peptide substrate; ^c^ Surface Plasmon Resonance (SPR) with PRMT5:MEP50 complex (K_d_) ^d^ Symmetric dimethylation in cell homogenates from Z-138 cells (EC_50_) ^e^ In-Cell Western (ICW): sDMA Western, various cell lines (IC_50_); ^f^ Symmetric arginine methylation of SmD3 in Z-138 cells (IC_50_); ^g^ Symmetric dimethylation of SmBB’ in MCF7 cells (IC_50_); ^h^ Long Term Proliferation (LTP) assay, various cell lines (IC_50_); ^i^ Proliferation assay in Z-138 cells, (IC_50_); ^j^ Cancer cell line proliferation assay (IC_50_); n. a., not available. Literature for GSK3326595: Gerhart et al.,^*32*^ GSK3203591: Duncan et al. and Scheer et al.,^*28–30*^ LLY-283: Bonday et al.,^*14*^.

| Assay | EC<sub>50</sub> (μM) |  |  |
| --- | --- | --- | --- |
|  | GSK3326595 | GSK3203591 | LLY-283 |
| Biochemical | 0.0062 ± 0.0008 <sup>a</sup> | 0.004 - 0.011 <sup>b</sup> | 0.022 ± 0.003 <sup>b</sup> |
| SPR | n. a. | n. a. | 0.006 ± 0.002 <sup>c</sup> |
| ELISA | 0.0025 <sup>d</sup> | 0.0021 <sup>d</sup> | n. a. |
| ICW | 0.0003–0.014 <sup>e</sup> | 0.003–0.056 <sup>f</sup> | 0.025 ± 0.001 <sup>g</sup> |
| Proliferation | 0.0057–0.106 <sup>h</sup> | 0.062 <sup>i</sup> | 0.003–2.5 <sup>j</sup> |
| NanoBRET intact cell | 0.062 ± 0.004 | 0.019 ± 0.004 | 0.3 ± 0.1 |

### SAM influence on PRMT5 inhibitor target engagement

To simulate lower SAM concentration levels as they might occur in other cell types *in vivo*, we performed TE assays using CBH-002 in cell lysates. Due to the dilution effect, these experiments yielded EC_50_ values closer to the biochemical results reported in case of LLY-283 (Fig. 3A). Notably, the decreased SAM concentration in the cell lysates had exactly the opposite effect on the substrate-mimetic inhibitors GSK3326595 and GSK3203591 with their respective EC_50_ values increasing by approximately 100-fold (Fig. 3A). For the structurally related inhibitor GSK3235025 (EPZ015666), previous mechanistic studies suggested that presence of SAM in the active site may potentially augment PRMT5 binding affinity by helping to stabilise inhibitor binding in the substrate pocket.^*15*^

**Figure 3:**
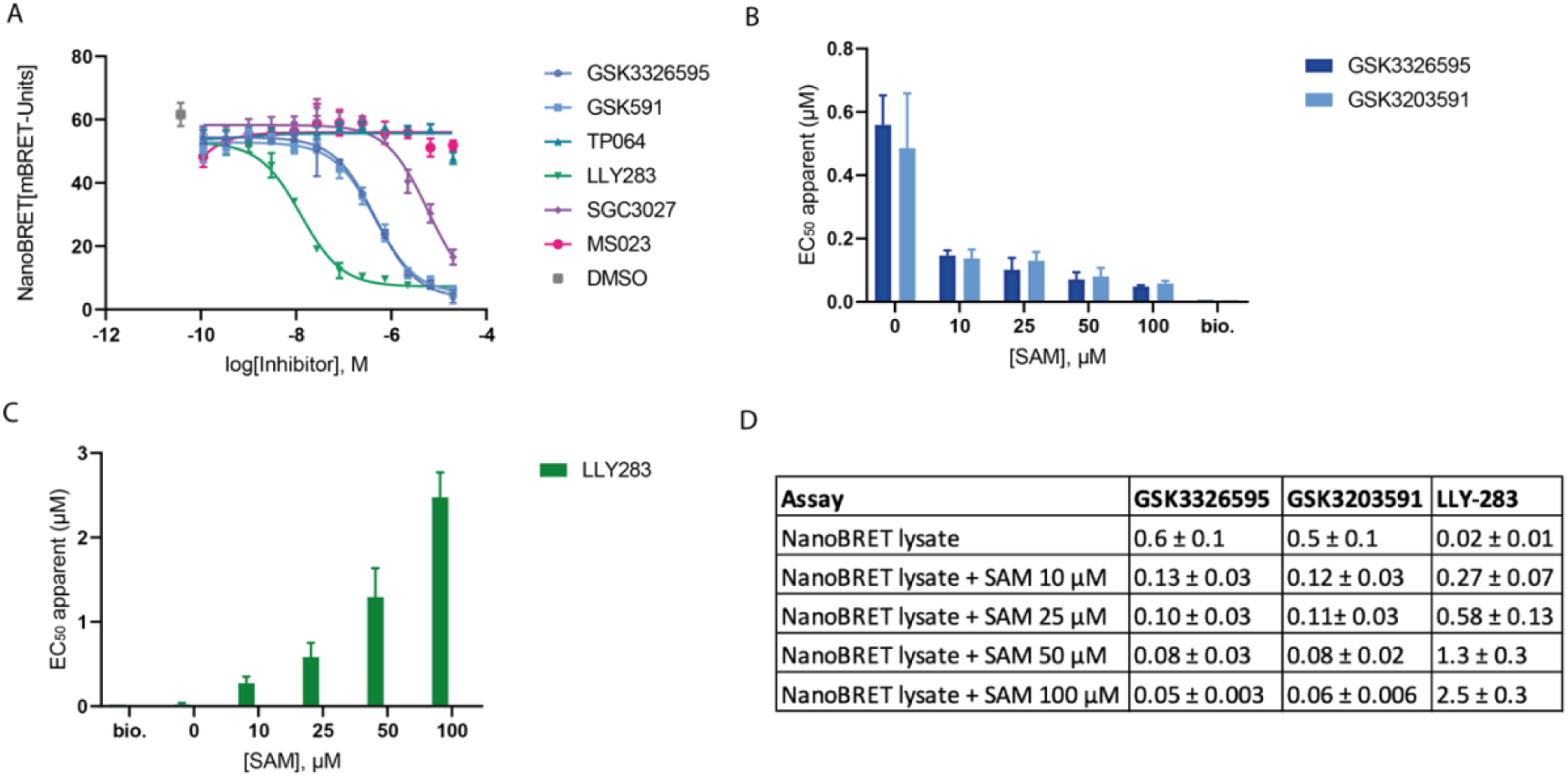
Effect of SAM levels on cellular EC_50_ values of PRMT5 inhibitors. A) HEK293 cells were lysed before plating and then treated as stated in Fig. 2. B) and C) Influence of different SAM concentrations on compound EC_50_ values in lysates (bio. = representative biochemical IC_50_ value from literature, see Table 1). D) Summary table for SAM lysate experiments, EC_50_ values in μM.

To further investigate this phenomenon, we next artificially elevated the SAM concentration in the lysates by exogenous addition of SAM at final assay concentrations of 10, 20, 50, and 100 μM, respectively. For LLY-283, previous reports suggested a non-competitive mechanism of action with regard to SAM.^*14*^ However, in our hands EC_50_ values were gradually increasing indicating a direct relationship with SAM levels and thus cofactor-competitive behaviour (Fig. 3C D,). Conversely, EC_50_ values for GSK3326595 and GSK3203591 (Fig. 3B, D) decreased upon SAM addition, further supporting the notion that cofactor complementation facilitates substrate-competitive inhibitor binding to PRMT5 via the formation of the ternary PRMT5:SAM:inhibitor complex as previously hypothesised.^*15*^

To further rationalise this concept, we re-evaluated the crystal structures of PRMT5 in complex with SAM as well as SAM- or substrate-mimetic inhibitors. In case of the substrate mimetic GSK3235025 (EPZ015666) the tetrahydroisoquinoline scaffold plays a significant role upon binding to PRMT5 (Fig. 4A). The bicyclic ring structure interacts with F327 via π-π stacking and takes advantage of a cation-π interaction with the positively charged sulfonium group of SAM.^*15*^ In absence of the inhibitor, SAM maintains the interactions with PRMT5 mainly via the adenosine core whereas the other part of the cofactor extends towards the substrate binding pocket. However, when GSK3235025 is bound, SAM adopts a different conformation to accommodate the inhibitor (Fig. 4B). Notably, F327 appears to assume distinct positions depending on whether SAM, GSK3235025 or LLY-283 are bound which likely explains our observation that CBH-002 is able quantify TE for both inhibitor types (Fig. 4B, C).

**Figure 4:**
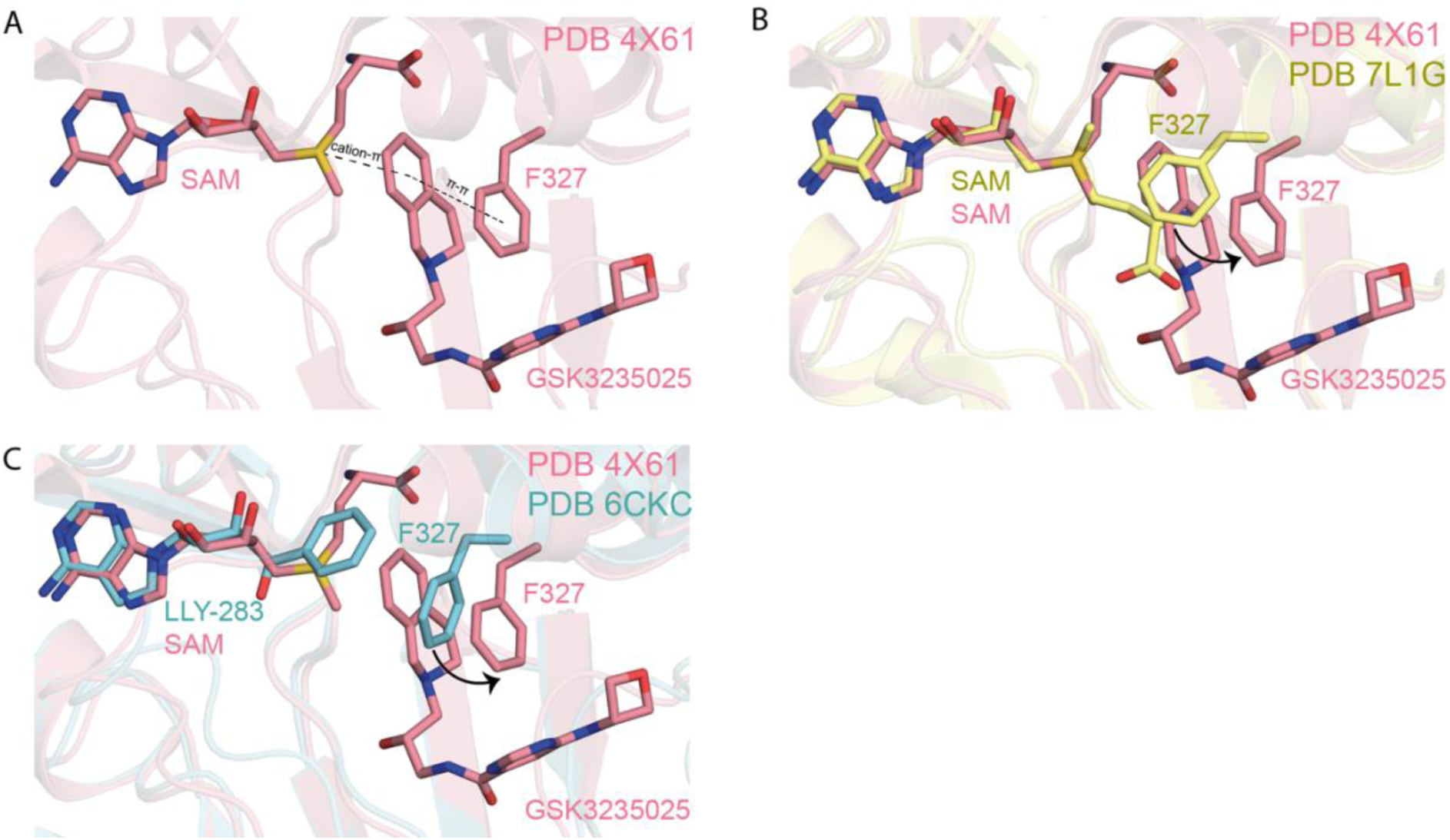
Co-crystal structures indicate mutually exclusive binding of SAM, LLY-283 and GSK3235025 (EPZ015666). A) SAM and GSK3235025 bind in a stacked manner (PDB: 4X61, pink). B) Induced conformational change of F327 when SAM or GSK3235025 are bound, respectively (PDB: 7L1G, yellow and 4X61, pink). C) Compared to the SAM-mimetic inhibitor LLY-283, binding of GSK3235025 requires re-positioning of F327 to avoid a clash with the inhibitor (PDB: 4X61, pink and 6CKC, blue).

### Proteome-wide selectivity profiling of GSK3326595

Having investigated the specific on-target occupancy of PRMT5 inhibitors, we next decided to evaluate the proteome-wide selectivity of the clinical candidate GSK3326595. To this end, we immobilised the amine-functionalised analogue CBH-001 on sepharose beads to generate an affinity matrix enabling chemoproteomic profiling.^*33*^ In order to distinguish genuine protein targets from non-specific binders we performed competitive binding experiments in KMS11 lysates which were pre-incubated with either DMSO or an excess of free GSK3326595 before analysis of matrix-enriched proteins. Western blot analysis confirmed potent enrichment of PRMT5 and its complex partner WDR77 (MEP50) which were efficiently competed by unmodified GSK3326595 (Fig. 5A and Supp. Fig. 2A). Subsequent mass spectrometry analysis suggested GSK3326595 to be highly specific for its cognate target with significant enrichment and competition only observed for PRMT5 alongside known interactors (Fig. 5B). For example, CLNS1A (plCln) has been found to act as a substrate adaptor for the PRMT5 complex.^*34*^ We also identified the zinc finger protein WIZ as significantly enriched in our CBH-001 matrix pull-downs. WIZ has been found to interact with PRMT5 in a large scale AP-MS study in HeLa cells^*35*^ and constitutes a core subunit of the protein methyltransferase G9a/GLP complex which catalyses histone methylation for transcription repression.^*36, 37*^ A di-methylated form of WIZ has been identified by MS analysis after enrichment of lysine and arginine methylated peptides in HeLa-S3 cells.^*38*^ These data in conjunction with the fact that WIZ contains a GRG motif which is preferred by PRMT5 for arginine methylation^*3*^ suggest WIZ as a candidate substrate of PRMT5. EPB41 is a member of the protein band 4.1 R superfamily of mammalian erythrocyte cytoskeletal proteins and has been shown to interact with the PRMT5 adaptor CLNS1A in a yeast-two-hybrid assay.^*39*^ Interestingly, the Rho GEF FARP1, which we also observed in the CBH-001 pull-down, is part of the same superfamily and shares the prominent FERM motif.^*40*^ FARP1 has not previously been reported as a PRMT5 complex binding partner but FARP2 has been identified as a PRMT5 interactor in AP-MS studies using HEK293T cells.^*41*^

**Figure 5:**
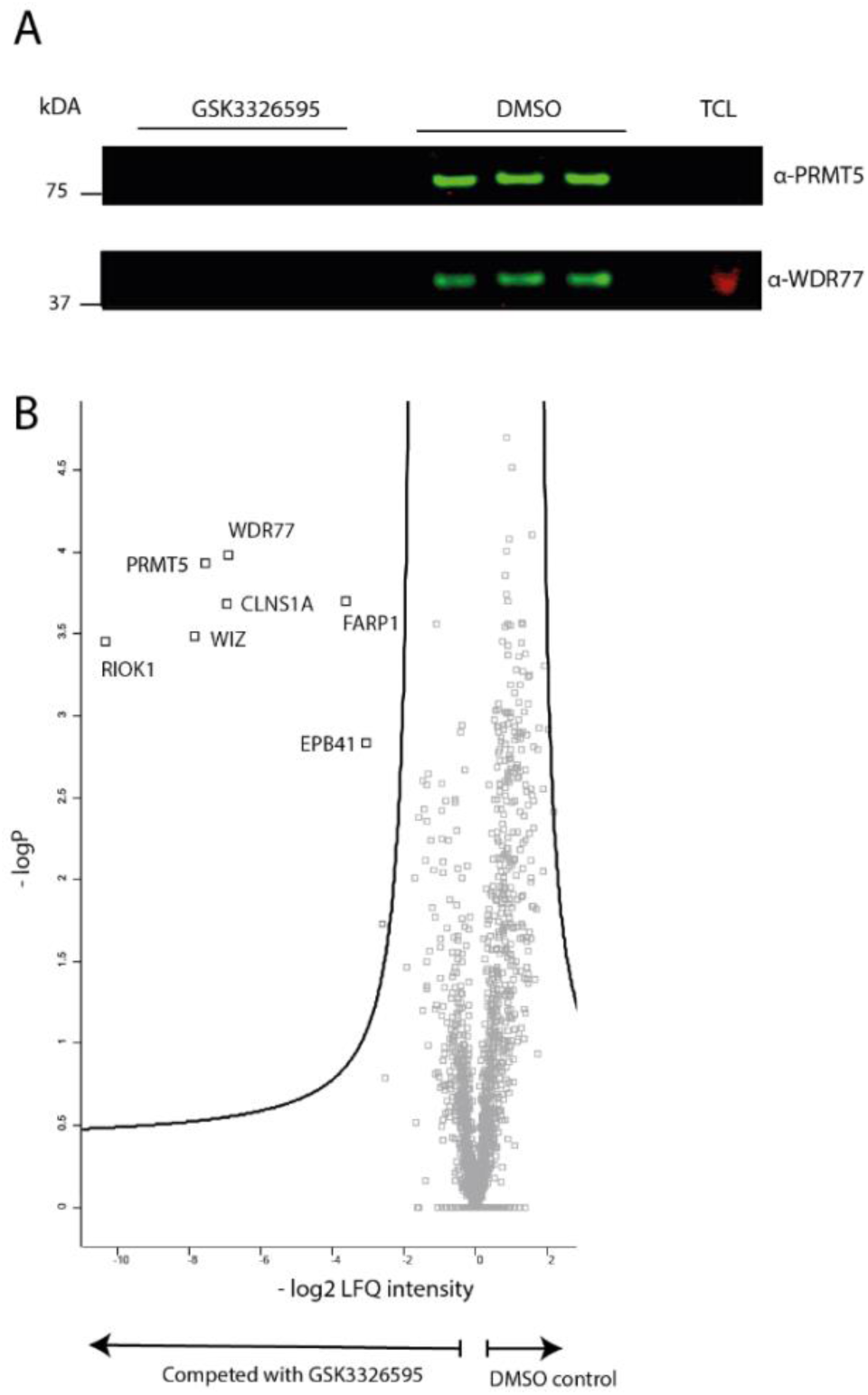
Proteome-wide specificity of GSK3326595 as determined by chemical proteomics. A) Western blot analysis of competitive PRMT5 engagement by affinity probe CBH-001. Competition with either parent inhibitor or DMSO in KMS11 lysate shows inhibitor-dependent enrichment of PRMT5 (72 kDa, green) and WDR77 (36 kDa, green) (TCL, total cell lysate). Red band indicates GAPDH which was used as TCL loading control. B) Volcano plot of CBH-001-enriched proteome from KMS11 cell lysate, with targets significantly competed by 20 μM GSK3326595 versus DMSO control (FDR = 0.05, S0 = 0.2).

To account for any potential bias introduced by installation of a linker moiety we also conducted orthogonal thermal profiling experiments with GSK3326595 in live KMS11 cells. Using a 2D setup,^*42*^ cells were treated with four different inhibitor concentrations (i.e. 5, 1, 0.2 and 0.04 μM) or DMSO, and subjected to a denaturing temperature gradient (42-64 °C). This approach allows for visualising dose-dependent thermal stabilisation of proteins, thereby increasing confidence in direct drug-target interactions. Following mass spectrometry analysis, 5,926 proteins were identified out of which 5,434 were detected with high FDR confidence. Data were processed with an R-script,^*42*^ as well as a bespoke, in-house coarse MATLAB filter we established for fast thermal profiling data deconvolution (Fig. 6A and Supp. Fig. 3). This analysis yielded two high confidence hits, the cognate target PRMT5 and its well-known binding partner WDR77. GSK3326595-dependent stabilisation of PRMT5 and WDR77 was also validated by Western blot (Figure 6B and Supp. Fig. 2B). Taken together, these results suggest that GSK3326595 is a highly specific PRMT5 inhibitor.

**Figure 6:**
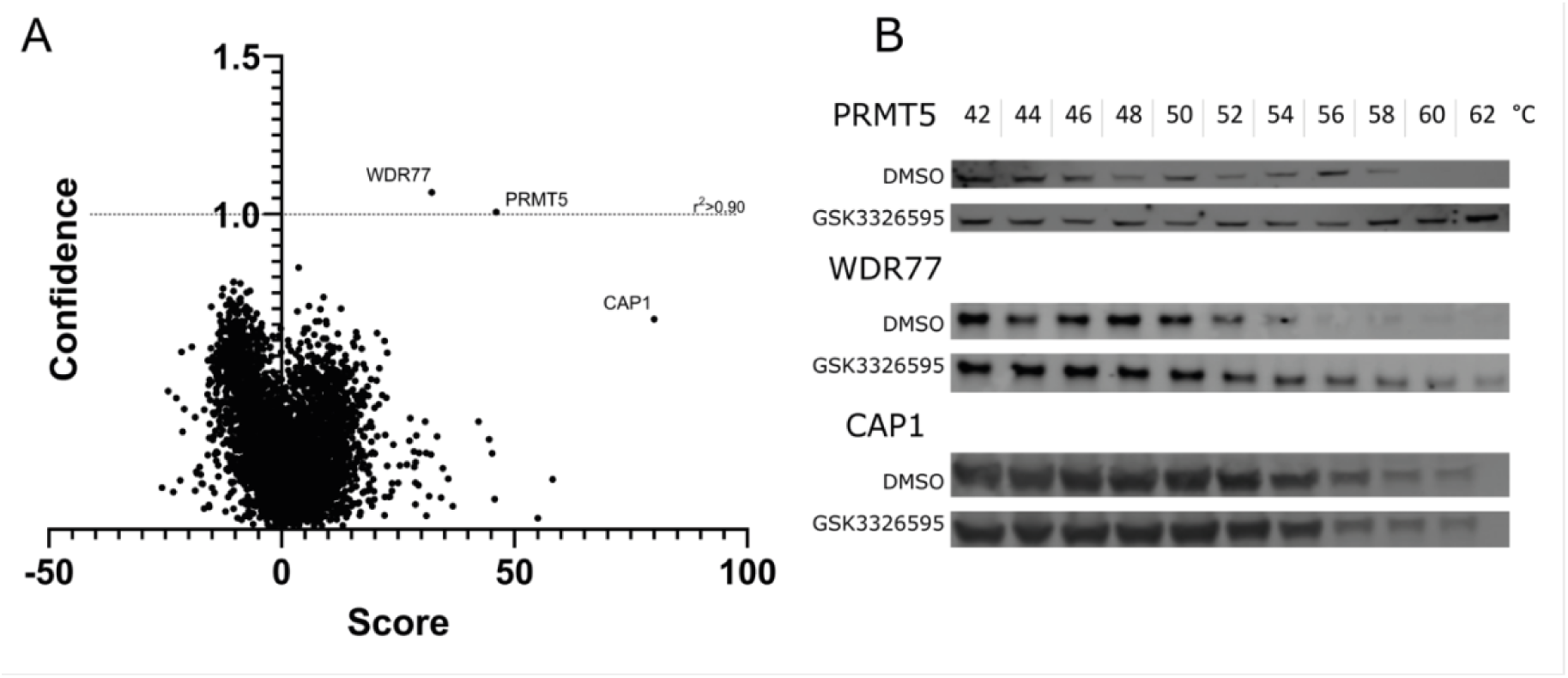
2D thermal profiling results for GSK3326595 identify the methylosome complex comprising PRMT5 and WDR77. A) Volcano plot. For each protein, the score indicates the volume under the surface of stabilization as a 2-dimensional function of temperature and compound concentration as calculated by the algorithm, and confidence is –log(1-r^2^) of the fitted surface (for more information about the algorithm, see Supplementary Figure 3). B) Validation of PRMT5 and WDR77 via Western blot CETSA using 5 μM GSK3326595.

## Discussion

PRMT5 inhibitors are advancing in clinical trials including tumours with high unmet clinical need such as multiple myeloma and glioblastoma ^*43, 44*^. Several studies have underscored the importance of correlating biochemical and cellular TE with functional pharmacological assays in drug development programmes to support biomarker identification and patient stratification.^*16, 17*^ Previously, cell TE for PRMT5 inhibitors has been demonstrated via imaging-based detection of symmetrically dimethylated nuclear proteins^*45, 46*^ as well as in a PRMT5-RIOK1 protein-protein interaction NanoBiT assay in permeabilised cells.^*47, 48*^ Recently, studies have shown that NanoBRET assays using bespoke ETP probes are a powerful means to study direct TE in a quantitative fashion ^*22, 49*^. While CETSA has been established for PRMT5 previously,^*15*^ a key advantage of the NanoBRET system is its ability to provide a continuous quantitative readout of cellular TE. Our intact cell results provide EC_50_ values that correlate with biochemically derived data and suggest an expected decrease in potency likely due to effects relating to cellular uptake, compound distribution and a more complex intracellular environment in general. In addition, by comparing live cell, lysate and SAM complementation experiments, we demonstrate differential cofactor effects on inhibitor binding expanding on previous observations made with purified recombinant proteins.^*15*^ In lysates, EC_50_ values for substrate-competitive inhibitors such as GSK3326595 are increased as SAM is diluted and therefore the cofactor cannot contribute to the formation of a favourable ternary complex. In case of GSK3326595, earlier mechanistic work showed that increasing pre-incubation times to 60 min in presence of SAM led to a reduction of the respective biochemical EC_50_ values in a PRMT5:MEP50 activity assay, suggesting a slow binding inhibition mode of action.^*32*^ Artificial addition of SAM facilitates substrate-mimetic inhibitor binding as indicated by more efficient displacement of CBH-002. It is worth noting that although GSK3326595 was marginally less potent than GSK3203591 in the intact cell NanoBRET, its EC_50_ values appeared to respond slightly better to exogenous addition of SAM, particularly at high cofactor concentrations. Conversely, LLY-283 which targets the SAM-pocket of PRMT5, performed better in lysate whereas increased SAM levels resulted in higher EC_50_ values, indicating cofactor-competitive behaviour. With regard to previously published data our results are largely consistent, the fact that our NanoBRET system uses HEK293 cells limits the ability to perform direct comparisons as this cell line has not been evaluated in the literature with these inhibitors. Notably, our findings reinforce the concept of SAM contributing to inhibitor binding which has also been demonstrated for PRMD9 using SPR.^*50*^ Also, the influence of MTAP deletion leading to higher MTA levels might as well influence inhibitor potency in cells.^*51*^

Intrigued by the ability of our substrate-mimetic ETP CBH-002 being able to survey both substrate- as well as cofactor-competitive PRMT5 inhibitors we re-examined previously published crystal structures to rationalise our observations. Our analysis strongly supports the idea that F327 plays a key role as a gatekeeper-like sensor which repositions in response to SAM- as well as substrate-mimetic inhibitors.^*52*^ Our ETP is able to exploit these dynamic interactions allowing for the detection of both substrate- as well as cofactor-mimetic inhibitors. We speculate that our assay could also be used to evaluate dual-competitive orthosteric and allosteric inhibitors.^*45, 46*^ A recent study found that spirodiamines based on a purine scaffold can occupy both the cofactor- as well as parts of the substrate-binding site.^*53*^ However, these compounds are SAM-competitive but non-competitive with regard to substrates. Interestingly, PRMT5 has a several accessible cysteine residues (C278, C449) which has enabled the development of covalent inhibitors.^*48, 52*^

Currently, there is limited data with regard to the global specificity of PRMT inhibitors. We have previously investigated the proteome-wide selectivity of the pan-type I inhibitor MS023.^*30*^ Here, we extend this effort to the clinical PRMT5 inhibitor GSK3326595 (pemrametostat) using complementary chemoproteomic and 2D thermal profiling experiments. We selected KMS11 as a model system since PRMT5 is overexpressed in multiple myeloma (MM) and has been suggested as a prognostic marker. Genetic knockdown and PRMT5 inhibition using GSK3235025 (EPZ015666) have been shown to inhibit growth and induce apoptosis in MM cell lines.^*44*^ Both experiments suggest remarkable selectivity of GSK3326595 for PRMT5, although there may be additional targets that are missed by MS detection or are not expressed in KMS11 cells. Nevertheless, a recently reported VHL-based PRMT5 PROTAC derived from GSK3235025 (EPZ015666) was found to be highly specific.^*54*^ Notably, we also identified several known PRMT5 interactors such as WDR77 (MEP50), RIOK1 and CLNS1A in our MS data. RIOK1 and CLNS1A have previously been found to bind PRMT5 in a mutually exclusive fashion.^*55*^ Our results therefore indicate that GSK3326595 can likely recognise both complexes. In this context, it is interesting to note that COPR5 is thought to utilise the same binding site as RIOK1 and CLNS1A^*5, 47*^ but was not detected in our experiments. However, it is possible that this complex is not stable under our experimental conditions or does not occur in KMS11 cells. With respect to the other proteins identified in our pull-down, FARP1 which was enriched and competed may be a potentially novel PRMT5 binding partner in human cells. Previous work did not detect any changes in methylated peptides for FARP1 upon GSK3203591 treatment in HeLa cells suggesting it may not be a substrate for PRMT5.^*3*^ In contrast, WIZ which contains a sequence motif preferred by PRMT5 for methylation and which was found to interact with PRMT5 in AP-MS studies could be a direct substrate. Pharmacologic inhibition of PRMT5 has been shown to result in a minor down-regulation of methylated EPB41^*3*^, which is known to bind CLNS1A and was co-purified in our drug pull-down. EPB41 has been linked to anaemia,^*56*^ yet any functional consequences of a this interaction remain to be elucidated.

In conclusion, our results reveal new insights into the mechanism of action of PRMT5 inhibitors and provide a set of tools which we envision to support the development of future inhibitors. Our proteomic profiling data highlight the selectivity of clinical PRMT5 inhibitors and how these compounds interact with PRMT5 adaptor complexes. The highly versatile NanoBRET system allows to correlate cofactor-dependent cell TE with functional readouts and can be easily expanded towards other PRMTs.

## Methods

All solvents were purchased of HPLC grade from commercial suppliers (Sigma, Thermo Fisher) and used without further purification. Anhydrous solvents were purchased from Acros Organics and stored under a nitrogen atmosphere with activated molecular sieves. CBH-001 was purchased from Enamine. NanoBRET 590 SE, NanoBRET™ Nano-Glo® Detection System and CellTiter-Glo® Luminescent Cell Viability Assay was purchased from Promega.

Reaction progress was monitored by TLC (Thin Layer Chromatography) and LC-MS. For the silica gel TLC, aluminium plates coated with 0.25 mm 60F_254_ silica gel (Merck) were used and visualized with UV light at l = 254 nm or l = 365 nm.

1H NMR spectra were recorded using a Bruker Avance 400 MHz spectrometer (400 MHz) and the deuterated solvent stated. Chemical shifts (δ) are quoted in parts per million (ppm) and referenced to the residual solvent peak. Multiplicities are denoted as singlet (s), doublet (d), triplet (t), quartet (q) and quintet (p) and derivatives thereof and multiplets (m). (br) denotes a broad resonance peak. Coupling constants are recorded as Hz and rounded to the nearest 0.1 Hz. Spectra were analysed using MestreNova 14.2.1.

LC-MS chromatograms determining mass and purity were obtained using a Waters Auto purification System equipped with Waters 2489 UV/Vis or Water 2998 Photodiode Array detectors, Waters 2424 ELS detector and SQ Detector 2 or Acquity QDa mass detector. A Phenomenex Kinetex 5μm EVO C18 100A, 3 × 100 mm column was used for analytical measurements and a Phenomenex 5 μm EVO C18 100 A, 21.2 × 150 mm column was used for preparative HPLC purifications using a gradient program (eluent I: acetonitrile/water = 5/95 with 20 mM ammonium acetate buffer, pH 6.0; eluent II: acetonitrile/water = 80/20 with 20 mM ammonium acetate buffer, pH 6.0).

Compound names were generated using ChemBioDraw Ultra v19.0 systematic naming.

### NanoBRET TE assay with CBH-002

Cells were split when reaching 80 % confluency. Each concentration was prepared in triplicates. HEK293 cells were prepared as 2 × 10^5^ cells/mL in DMEM (Dulbecco s Modified Eagle Media, Life Technologies) + 10% FBS (Foetal Bovine Serum, Life Technologies) medium and transfected with NL-PRMT5 (Promega) according to the manufacturer’s protocol using a transfection solution consisting of 1 μg DNA, 9 μg Transfection Carrier DNA (Promega) and 30 μL FuGENE HD Transfection Reagent (Promega) per 1 mL Opti-MEM (Life Technologies). After 10 min incubation at room temperature, the DNA:FuGENE complex was formed and the cells were transfected 1:20 with the transfection solution and subsequently incubated at 37 °C in a humidified 5 % CO_2_ atmosphere for 24 h.

Depending on the size of the experiment, either 6-well plates (Greiner-Bio) or T75 flasks (Greiner-Bio) were used for transfection. The BRET assay was performed in a 384-well plate (UltraCruz, PP) by plating 2 × 10^5^ cells/mL in OptiMEM; for lysates, Passive Lysis 5x buffer (Promega) was used and SAM (Sigma) was optionally added for competition experiments. The ETP was prepared as a 100x stock in 100 % DMSO (Sigma) and 20x stock in NanoBRET Tracer Dilution buffer (Promega) in a PP 384-well plate, to prevent interactions with the plate material. The test compounds (GSK3203591 (Sigma) GSK3326595, MS023, LLY-283 and SCG3027 (MedChem Express), TP064 (Tocris)) were prepared as 1000x stock in 100 % DMSO and 10x stock in Opti-MEM in a 96-well plate (Starlab). As controls, no compound and no energy transfer probe/energy transfer probe wells were prepared. After addition of the energy transfer probe and the compounds to the wells, the plates were allowed to incubate at 37 °C and 5 % CO_2_ for 2 h. Next, the BRET measurement was prepared by adding a mix of NanoBRET Nano-Glo Substrate (1:166) and Extracellular NanoLuc Inhibitor (1:500) in Opti-MEM to the wells. Followed by 10 min incubation at room temperature, the plates were measured at the PheraSTAR FSX at 450 nm (donor) and 610 nm (acceptor).

For the data analysis to create raw BRET ratio values, the acceptor value was divided by the donor. For background correction, the no energy transfer probe ratio was subtracted from these values. Next, the corrected BRET ratio values were multiplied by 1000 to obtain milli BRET units (mBU) which are then plotted against the energy transfer probe.

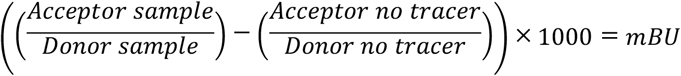

### Cell Titer Glo Luminescent Cell Viability Assay

Cell Titer Glo Luminescent Cell Viability Assay Kit (Promega) was used and subsequently performed after the BRET measurement. The components were mixed according to the manufacturer’s recommendation, diluted 1:3 with DPBS (Life Technologies) and 25μL added to the wells. After 10 min incubation at room temperature, the plates were measured at the PheraSTAR FSX. For the data analysis, the mean and SEM (Standard Error of the Mean) of the raw triplicate luminescence values were calculated. Subsequently, these values were normalized to the mean of the positive control (DMSO, 100 % viability), which is obtained from wells containing no compound and no energy transfer probe and plotted against the logarithm of the compound concentration.

### Pulldown

For profiling PRMT5 inhibitor GSK3326595, label-free quantification (LFQ) mass spectrometry was conducted. Chemical proteomics was conducted as previously described.^*30*^ In brief, KMS11 (ATCC) were cultured at 37 °C, in a humidified 5 % CO_2_ atmosphere in RPMI-1640 (Life Technologies) + 10 % FBS. To obtain lysate, the cells were harvested at 80 % confluency from 5x T75 flasks (Greiner) and subsequently pelleted and washed with DPBS. The obtained pellets were then lysed by addition of 3x pellet volume of lysis buffer (50 mM Tris pH 7.5, 0.8 % v/v NP-40, 5 % v/v glycerol, 1.5 mM MgCl_2_, 100 mM NaCl, 25 mM NaF, 1 mM Na_3_VO_4_, 1 mM PMSF, 1 mM DTT, 10 μg/mL TLCK, 1 μg/mL

Leupeptin, 1 μg/mL Aprotinin, 1 μg/mL soybean trypsin) on ice, before being pushed gently 10 × through a 21 G needle (Braun) using a 1 mL syringe (Braun, Inject-F). The lysates were allowed to incubate on ice for 10 min before being ultracentrifuged at 17000 G at 4 °C for 30 min. Next, total protein concentration was adjusted to 10 mg/mL and the lysates were stored for further use at −80 °C. In preparation for a pulldown, the amine derivatized compound CBH-001 was coupled to NHS-activated Sepharose 4 Fast Flow beads (Cytiva). 100 μL beads as a slurry in 50% isopropanol were used for each experiment, the experiments were done in triplicates (with/without competition). The beads were washed with 500 μL DMSO and centrifuged at 0.1 G at RT for 3 min. The supernatant was removed, and the washing step was repeated 3x. Then the bead bed was re-suspended in 50 μL DMSO. To immobilise the couplable compound on the beads, 2 μL of a 10mM stock in DMSO was added to the beads along with 0.75 μL DIPEA (Diisopropylethylamine). Then the tubes were incubated on the rotor shaker at 10 RPM for 20 h at RT. Subsequently, successful immobilisation and depletion of the free amine from the supernatant was determined by LC-MS analysis. The remaining unreacted NHS (N-Hydroxysuccinimide) groups were blocked with 2.5 μL ethanolamine and incubation again at the rotor shaker 10 RPM for 8 h at RT. The lysates were thawed on ice before pre-treatment with DMSO control or parent compound, GSK3326595, at 10mM and incubating for 30 min at 4 °C. A total cell lysate (TCL) sample was kept aside. The derivatized beads were washed 3x with 500 μL DMSO and then 3x with freshly prepared lysis buffer by centrifugation at 0.1 G for 3min at RT and subsequent removal of the supernatant. After that, the derivatized beads were combined with the cell lysate at 1.6 mg of protein per pull-down (10 mg/mL) being either compound or DMSO control pre-treated in triplicates. Beads and lysates were then incubated on the rotor shaker at 10 RPM in the cold room at 4 °C for 2 h. Then the samples were centrifuged at 0.1 G for 3 min at 4 °C and 300 μL of the supernatant was removed. Working in the cold room at 4 °C, the slurry was resuspended in lysis buffer and 500 μL were transferred to a Bio Spin column (Bio-Rad) and the beads were allowed to settle by gravity. Next, the columns were washed with 5 mL lysis buffer via gravity flow and centrifuged at 0.1 G for 1 min at 4 °C to the remove the remaining supernatant.

Then the protein was eluted for Western Blot experiments using 100 μL of 2x sample buffer (65.8 mM Tris-HCl pH 6.8, 26.3 % (w/v) glycerol, 2.0 % SDS, 0.01 % bromophenol blue, 50 mM DTT) and boiling the beads at 100 °C for 10 min. The samples were stored in the freezer at −20 °C.

### Western Blot validation of Pulldown

The eluted proteins were analysed using a polyacrylamide gel and 1x MES buffer (50 mM MES, 50 mM Tris Base, 0.1 % SDS, 1 mM EDTA, pH 7.3) and transferal to a nitrocellulose blotting membrane. The membrane was blocked with blocking buffer (2.5 % m/v) BLOT-Quick Blocker (Merck) in PBST (Phosphate-buffered saline with Tween: 4.3 mM Na_2_HPO_4_, 1.47 mM KH_2_PO_4_, 137 mM NaCl, 2.7 mM KCl, 0.05 % (v/v) Tween 20) on a shaker in the dark for 1 h at RT. Then the blot was probed with the antibodies EPR5772 (Abcam, ab109451, 1:10000), MEP50 (2823S, Cell Signaling Tech., 1:1000) and G-9 (s-365062, Santa Cruz, 1:200) and imaged at the Odyssey CLx (Li-cor) at wavelength 700 and 800.

### Mass spectrometry sample preparation

100 μL of each sample were combined 1:1 freshly prepared 0.1 M TRIS solution. Then 5 μL 200 mM DTT was added, and the mixture incubated for 1 h at RT and was subsequently alkylated with 20 μL 200 mM iodoacetamide and incubated for 30 min in the dark at RT. The samples were diluted to 300 μL with TEAB and incubated at 37 °C overnight. To precipitate the proteins, 600 μL MeOH and 150 μL Chloroform were added to each vial and after vortexing 450 μL MilliQ-water were added. After centrifugation at 14.8 RPM for 5 min at RT, the upper phase was gently pipetted off without touching the precipitate at the interface. Next, 450 μL of MeOH was added, and the upper phase was pipetted off gently again. This was repeated once and then the samples were centrifuged at top speed. The supernatant was removed the vials left to dry 60 min RT. The precipitate was resuspended in 50 μL 6M urea buffer, followed by vortexing and sonication for 5 min. The solution was diluted with 250 μL MilliQ-water. Then trypsin in a 1:50 ratio regarding the total protein content was added, followed by incubation at 37 °C overnight. A SEP-PAK C18 purification was performed using solution A (98 % MilliQ-water, 2 % Acetonitrile, 0.1 % Formic acid) and solution B (35 % MilliQ-water, 65 % Acetonitrile, 0.1 % Formic acid). The samples were acidified with 1 % formic acid. Sola HRP SPE cartridge (Thermo Fisher) were attached to a vacuum manifold. First, the columns were equilibrated with 500 μL solution B. Second, 1000 μL of solution A were put on the column and again a small supernatant was left to prevent the column from running dry. Third, the peptide digest sample was loaded on the column. After that, the column was washed again with 1000 μL solution A. The columns were placed after the washing in fresh 1.5 mL Eppendorf vials. Then 600 μL solution B was used to elute into a fresh vial. Then, the samples were dried in the speed vac for 24h. Mass spectrometry data were acquired at the Discovery Proteomics Facility (University of Oxford). Peptides were resuspended in 5% formic acid and 5% DMSO and then trapped on an Acclaim™ PepMap™ 100 C18 HPLC Columns (5μm x 0.1mm x 20mm, Thermo Fisher Scientific) using solvent A (0.1% Formic Acid in water) at a pressure of 60 bar and separated on an Ultimate 3000 UHPLC system (Thermo Fischer Scientific) coupled to a QExactive mass spectrometer (Thermo Fischer Scientific). The peptides were separated on an Easy Spray PepMap RSLC column (75μm i.d. x 2μm x 50mm, 100 Å, Thermo Fisher) and then electro sprayed directly into an QExactive mass spectrometer (Thermo Fisher Scientific) through an EASY-Spray nano-electrospray ion source (Thermo Fisher Scientific) using a linear gradient (length: 60 min, 5% to 35% solvent B (0.1 % formic acid in acetonitrile), flow rate: 250 nL/min). The raw data was acquired on the mass spectrometer in a data-dependent mode (DDA). Full scan MS spectra were acquired in the Orbitrap (scan range 380-1800 m/z, resolution 70000, AGC target 3e6, maximum injection time 100 ms). After the MS scans, the 15 most intense peaks were selected for HCD fragmentation at 28 % of normalised collision energy. HCD spectra were also acquired in the Orbitrap (resolution 17500, AGC target 1e^5^, maximum injection time 128 ms) with first fixed mass at 100 m/z.

### MS Data analysis

Raw data was processed using MaxQuant version 1.6.1.2 and the reference complete human proteome FASTA file (UniProt). Label-Free Quantification (LFQ) and Match Between Runs were selected; replicates were collated into parameter groups to ensure matching between replicates only. Cysteine carbamidomethylation was selected as a fixed modification and methionine oxidation as a variable modification. Default settings for identification and quantification were used. Specifically, a minimum peptide length of 7, a maximum of 2 missed cleavage sites, and a maximum of 3 labelled amino acids per peptide were employed. Through selection of the ‘trypsin/P’ general setting, peptide bond cleavage at arginine or lysine (followed by any amino acid) was considered during in silico digest of the reference proteome. The allowed precursor and fragment ion mass tolerances were 4.5 ppm and 20 ppm, respectively. Peptides and proteins were identified utilizing a 0.01 false discovery rate, with “Unique and razor peptides” mode selected for both identification and quantification of proteins (razor peptides are uniquely assigned to protein groups and not to individual proteins). At least 2 razor + unique peptides were required for valid quantification. Processed data was further analysed using Perseus version 1.6.2.1 and Microsoft Excel. Peptides categorized by MaxQuant as ‘potential contaminants’, ‘only identified by site’ or ‘reverse’ were filtered, and the LFQ intensities transformed by log2. Experimental replicates were grouped, and two valid LFQ values were required in at least one experimental group. Statistically significant competition was determined through the application of P2 tests, using a permutation-based FDR of 0.05 and an S0 of 2, and visualized in volcano plots. Significantly competed targets were further analysed in STRING (http://string-db.org) and protein interaction networks generated. Basic STRING settings were used for network analysis of enriched proteins. The network edges represent confidence in interaction. Line thickness indicates the strength of data support with a minimum required interaction score of 0.400. All active interaction sources (Text mining, Experiments, Databases, Co-expression, Neighbourhood, Gene Fusion, Co-occurrence) were considered.

### Thermal Profiling

2D thermal profiling was performed according to previously described protocols.^*42*^ KMS-11 cells were grown until confluent in T-175 flasks (Greiner). To five flasks of confluent KMS-11 cells GSK33265905 was added up to the concentrations of: 5, 1, 0.2, 0.04 μM or equivalent volume of DMSO for one hour. The cells were then detached using the Tryp-LE trypsin replacement enzyme (Gibco) and pelleted into 12 aliquots per group. Each aliquot was heated to different temperatures for 3 minutes in a PCR machine (Bio-Rad), with the temperature range being 42-64 °C and 2-degree intervals. Cell pellets were then lysed in 0.1% NP-40 Tris-NaCl lysis supplemented with protease and phosphatase inhibitors (Sigma). Cell lysis was facilitated by three cycles of rapid freeze-thawing in liquid nitrogen. The lysates were clarified by centrifugation at 17,000 × g for 20 minutes, and BCA assay was performed on the soluble fraction. For MS analysis, 100 μg of protein per aliquot was taken.

### MS Analysis of TP samples

To reduce and alkylate proteins, lysate was first incubated with DTT up to a final concentration of 5 mM for 1 hour at RT followed by a 1 h incubation with iodoacetamide added to a final concentration of 20 mM. The proteins were then acetone-precipitated overnight –20°C and pelleted at 8,000 x g for 10 minutes at 4 °C. Dry pellets were resuspended in 100 μL of 50 mM TEAB and 2.5 μg of Trypsin/LysC (Promega) was added for overnight digestion at 37 °C. TMT labelling (Thermo Fisher) was performed according to the manufacturer’s protocol, and the samples were pooled according to the following scheme:

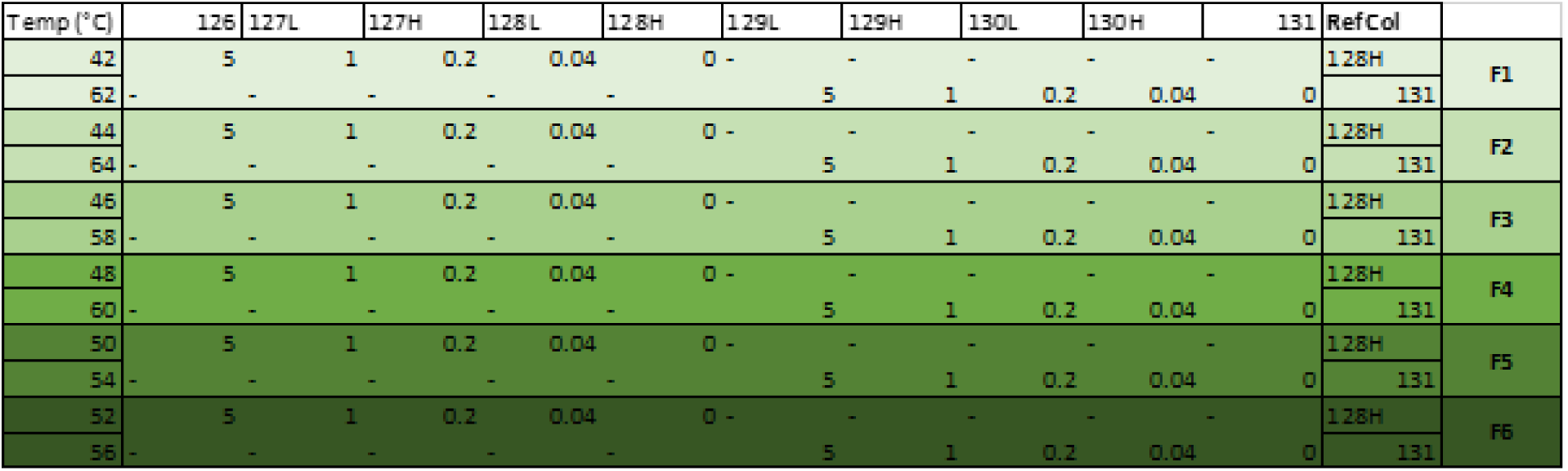

Pooled samples were subsequently desalted in C-18 columns (Pierce) and the solvent removed under reduced pressure in a SpeedVac Vacuum concentrator. Dry peptide mass was then resuspended in 120 μL of 98% MilliQ-H_2_O, 2% Acetonitrile, 0.1% Formic Acid (v/v) and fractionated using an UltiMate 3000 HPLC system (Thermo) at pH 10 using a 60 min gradient of 0% to 90% Acetonitrile in Water. Fractions were then dried using a SpeedVac Vacuum concentrator and resuspended in 100 μL of 98% MilliQ-H_2_O, 2% Acetonitrile, 0.1% Formic Acid (v/v). MS analysis was performed by nano-HPLC–MS/MS using a Dionex Ultimate 3000 nano HPLC with EASY spray column (75 μm × 500 mm, 2 μm particle size, Thermo Scientific) with a 60 min gradient of 2% to 35% (v/v) Acetonitrile in Water with 5% (v/v) DMSO and 0.1% (v/v) Formic Acid at a flow rate of 250 nL/min (600 bar per 40 °C column temperature). MS1 survey scans were acquired at a resolution of 60,000 at 375-1500 m/z and the 20 most abundant precursors were selected for CID fragmentation in a HCD cell. MS2 data were analysed in Thermo Proteome Discoverer 2.1 according to the manufacturer’s protocol.

### Western Blot CETSA

CETSA was performed according to previously reported procedures.^*21*^ KMS11 cells were grown until confluent in T-175 flasks (Greiner). One flask was treated with 5 μM GSK33265905 in DMSO, whereas the control flask was treated with an equivalent volume of DMSO for one hour. The cells were then harvested and aliquoted for heating to different temperatures for 3 minutes and lysed in NP-40 lysis buffer. The lysate was clarified by centrifugation at 17,000 × g for 20 minutes and 30 μg of protein was used for SDS-PAGE and Western Blotting. Antibody used: anti WDR77 (Cell Signaling, 2823), 1:100.

The mass spectrometry proteomics data have been deposited to the ProteomeXchange Consortium via the PRIDE partner repository with the dataset identifier PXD028138.

## Acknowledgements

EMR, JAW and KVMH are grateful for support by Myeloma UK and Bayer AG. This project has received funding from the Engineering and Physical Sciences Research Council (EPSRC) and the Medical Research Council (MRC) [grant number EP/L016044/1] as well as the Innovative Medicines Initiative 2 Joint Undertaking (JU) under grant agreement No 875510. The JU receives support from the European Union’s Horizon 2020 research and innovation programme and EFPIA and Ontario Institute for Cancer Research, Royal Institution for the Advancement of Learning McGill University, Kungliga Tekniska Hoegskolan, Diamond Light Source Limited. The authors would like to thank J. Bennett as well as B. Kessler, S. Bonham and R. Fischer from the TDI Discovery Proteomics Facility for their support and P. Brennan and all members of the Huber lab for valuable discussions about the project.

## Disclaimer

This communication reflects the views of the authors and the JU is not liable for any use that may be made of the information contained herein.

