## Supplemental Information for "A chemical biology toolbox to investigate in-cell target engagement and specificity of PRMT5-inhibitors"

**Supplement**

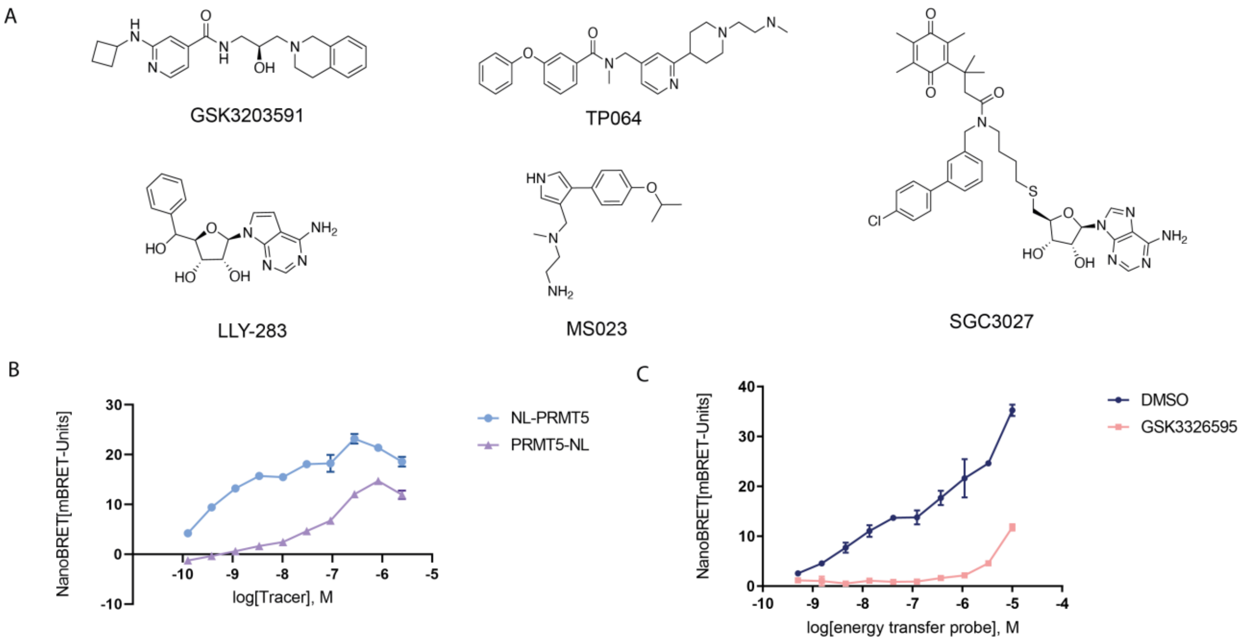


**Supplementary Figure 1:** Inhibitors used in this study and assay optimisation experiments. A) PRMT inhibitors tested in this study. B) Comparison of N- and C-tagged NLuc PRMT5 fusion proteins tested against 10 µM GSK3326595 or DMSO as a control. C) Titration of ETP CBH-002 against 10 µM inhibitor or DMSO.

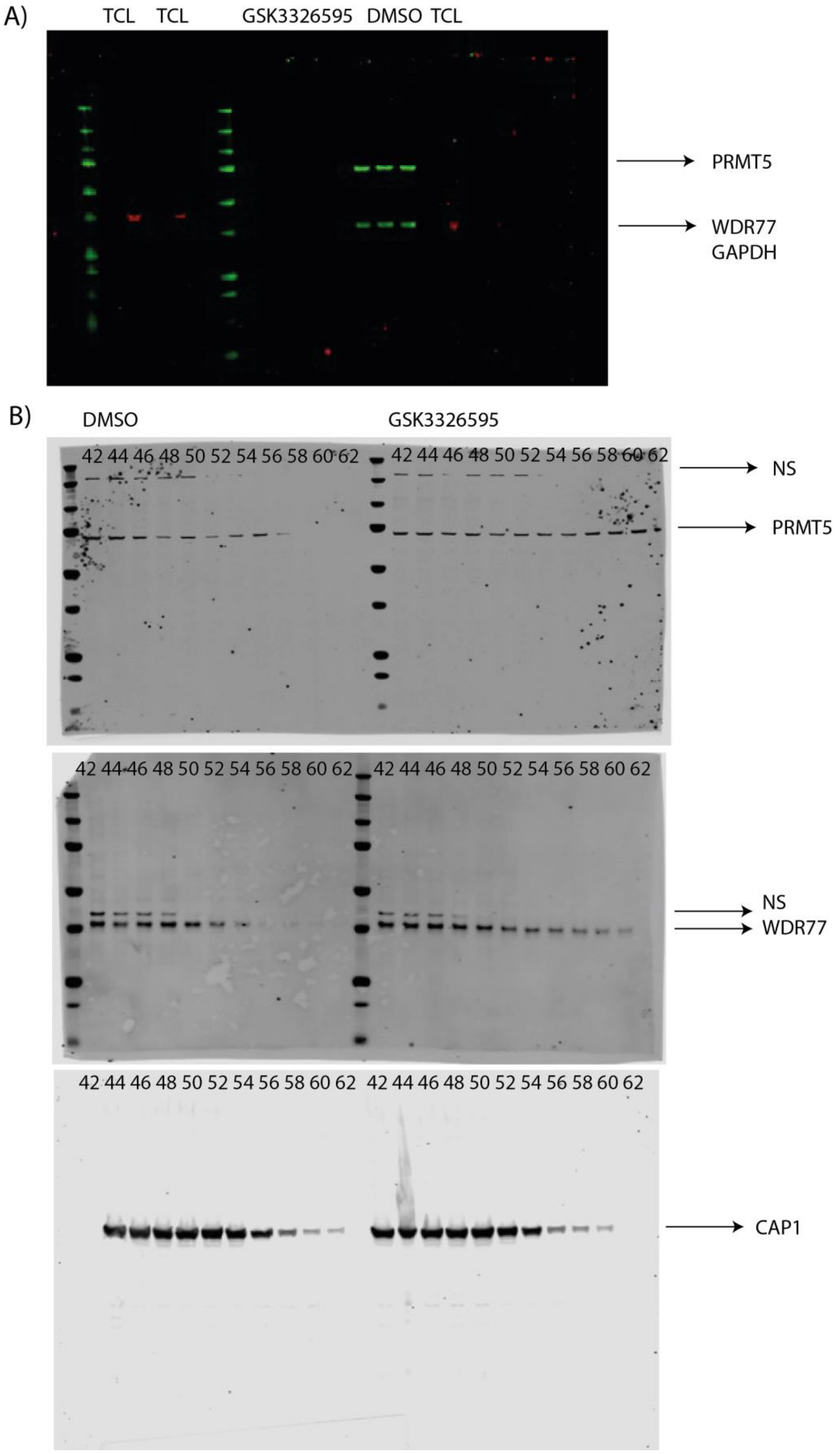
A)
TCL TCL GSK3326595 DMSO TCL
PRMT5
WDR77
GAPDH
B)
DMSO
GSK3326595
42 44 46 48 50 52 54 56 58 60 62 42 44 46 48 50 52 54 56 58 60 62
NS
PRMT5
42 44 46 48 50 52 54 56 58 60 62 42 44 46 48 50 52 54 56 58 60 62
NS
WDR77
42 44 46 48 50 52 54 56 58 60 62 42 44 46 48 50 52 54 56 58 60 62
CAP1

**Supplementary Figure 2**: Unmodified Western Blots. A) Western blot analysis of competitive PRMT5 engagement by affinity probe CBH-001. Competition with either parent inhibitor or DMSO in KMS11 lysate shows enrichment of PRMT5 (72 kDa, green) and WDR77 (36 kDa, green) (TCL, total cell lysate) (right) and including test concentrations (left). B) Unmodified Western Blot images depicting thermal stabilisation of putative GSK3326595 TPP hits ranging from 42-62 °C. Top: Stabilisation of PRMT5. Middle: Stabilisation of WDR77. Bottom: Stabilisation of CAP1. Note: due to exposure settings, the ladder signal was not intense enough to be visible; no other bands were observed. NS: Non-specific band.

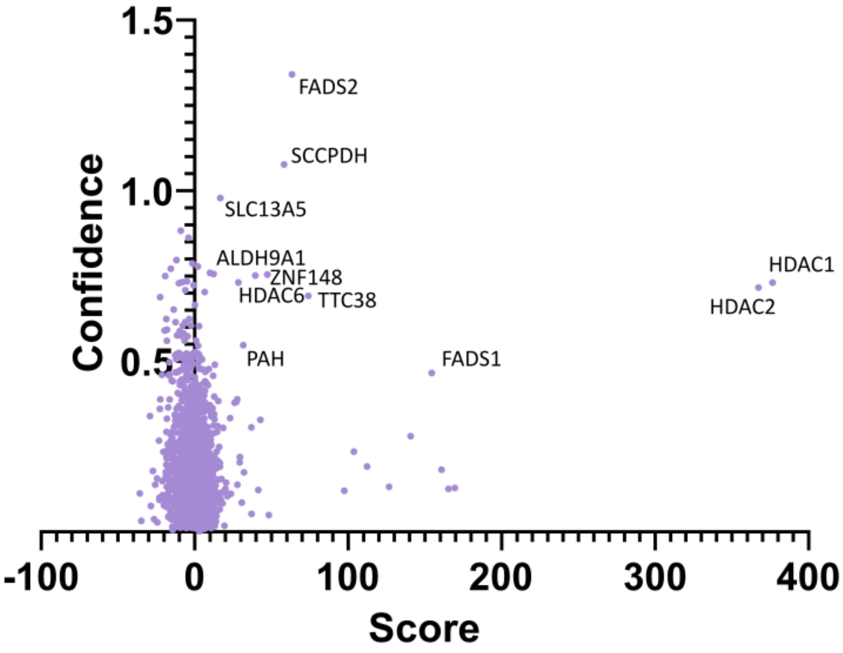


| 2D-TPP in cells: 15 stabilized >1 peptide, >1.5fold twice | |
|---|---|
| **clustername** | **IPI number** |
| ALDH9A1 | IPI00479877 |
| BAT5 | IPI00791049 |
| EPHX2 | IPI00104341 |
| **FADS1** | **IPI00784651** |
| **FADS2** | **IPI00183786** |
| FMO5 | IPI00215760 |
| **HDAC1** | **IPI00013774** |
| **HDAC2** | **IPI00289601** |
| **HDAC6** | **IPI00005711** |
| MIER1 | IPI00477825 |
| **PAH** | **IPI00017579** |
| SCCPDH | IPI00329600 |
| TBP | IPI00022831 |
| **TTC38** | **IPI00550644** |
| ZNF148 | IPI00010833 |

**Supplementary Figure 3:** Validation of the coarse MATLAB hit selector on the Becher 2016 dataset. As the output of the Becher TPP R-script produces essentially a 5x12 matrix for upwards of 6000 different proteins, it became necessary to develop an algorithm that would, at least coarsely, identify potential hits and reject false positives. The MATLAB filter treats each matrix as a 2D matrix and approximates the sigmoidal relationships between compound concentration and protein abundance with a linear function while simultaneously approximating the relationship between temperature and protein abundance with a polynomial equation. The resulting 2D surface equation is fitted over the data points, and the volume under the surface represents the magnitude of stabilisation, while the goodness of fit (r2) is a measure of confidence in the data. The algorithm has successfully been used to identify the hits from the Becher et al dataset. Note that the algorithm is only a coarse screen, and a manual inspection of the data is recommended.

**Synthesis**

**(*S*)-6-((1-(5-(3-(5,5-difluoro-7-(1H-pyrrol-2-yl)-5H-5λ4,6λ4-dipyrrolo[1,2-c:2',1'-f][1,3,2]diazaborinin-3-yl)propanamido)pentanoyl)piperidin-4-yl)amino)-N-(3-(3,4-dihydroisoquinolin-2(1H)-yl)-2-hydroxypropyl)pyrimidine-4-carboxamide (CBH-002)**.

(*S*)-6-((1-(5-aminopentanoyl)piperidin-4-yl)amino)-*N*-(3-(3,4-dihydroisoquinolin-2(1*H*)-yl)-2-hydroxypropyl)pyrimidine-4-carboxamide (5.4 mg, 0.010 mmol) was dissolved in 1.0 mL of anhydrous DMF. To the stirred mixture DIPEA (5.6 µL, 0.030 mmol) was added and stirring was continued for 10 min. To the clear colourless solution, NanoBRET® 590 SE (5 mg, 0.012 mmol) was added and the reaction was stirred to completion in the dark for 2 h. The sample was dried overnight *in vacuo*. The crude residue was re-solved in DMSO and subjected to reverse-phase preparative HPLC purification. Product containing fractions were concentrated *in vacuo*, affording the product (6.0 mg, 0.007 mmol, 69.0 %) as a purple solid. The purity was determined by LC/MS 98.2 %. LC/MS (ESI-1) found 819.4 g/mol (820.45 g/mol calculated for $C_{44}H_{55}BF_2N_{10}O_3$). 1H NMR (400 MHz, DMSO) δ 11.41 (s, 1H), 8.74 (d, J = 6.0 Hz, 1H), 8.29 (d, J = 1.2 Hz, 1H), 7.92 (t, J = 5.5 Hz, 1H), 7.76 (d, J = 7.6 Hz, 1H), 7.44 (s, 1H), 7.37 (s, 1H), 7.34 (d, J = 4.5 Hz, 1H), 7.28 (s, 1H), 7.17 (d, J = 4.6 Hz, 1H), 7.12 – 7.09 (m, 3H), 7.06 (d, J = 1.2 Hz, 1H), 7.02 (s, 1H), 6.33 (d, J = 4.0 Hz, 1H), 4.97 (d, J = 4.7 Hz, 1H), 4.23 (s, 2H), 4.08 (s, 1H), 3.92 – 3.80 (m, 3H), 3.61 (d, J = 5.9 Hz, 3H), 3.18 – 3.05 (m, 4H), 2.86 – 2.64 (m, 4H), 2.38 – 2.28 (m, 1H), 1.90 (s, 1H), 1.53 – 1.38 (m, 5H), 1.36 (s, 6H), 1.24 (s, 1H).

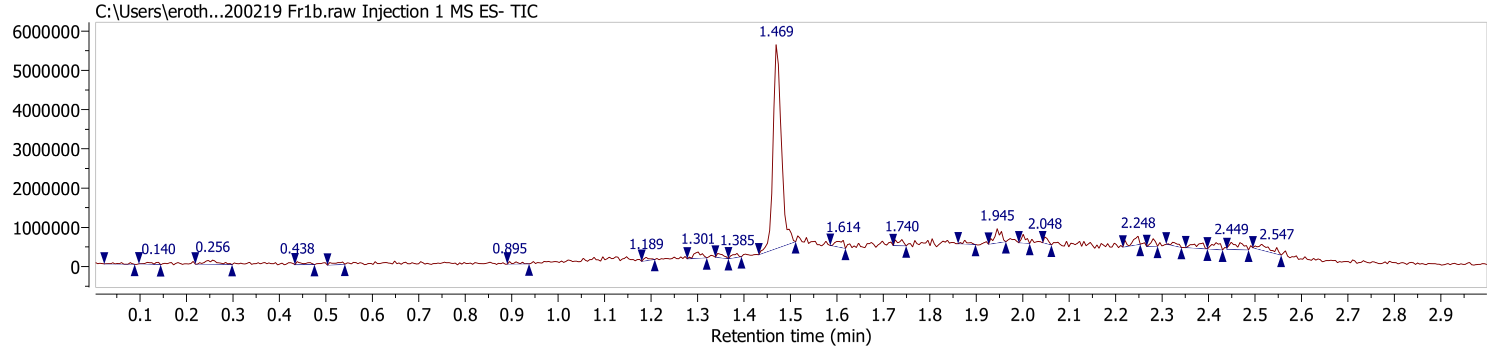
C:\Users\eroth...200219 Fr1b.raw Injection 1 MS ES- TIC
1.469
0.140
0.256
0.438
0.895
1.189
1.301
1.385
1.614
1.740
1.945
2.048
2.248
2.449
2.547
Retention time (min)

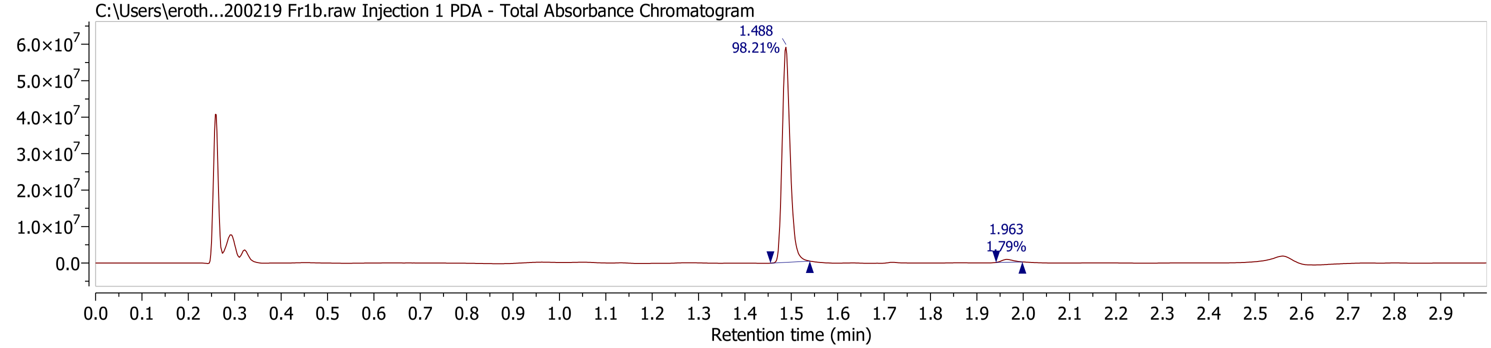
C:\Users\eroth...200219 Fr1b.raw Injection 1 PDA - Total Absorbance Chromatogram
1.488
98.21%
1.963
1.79%
Retention time (min)

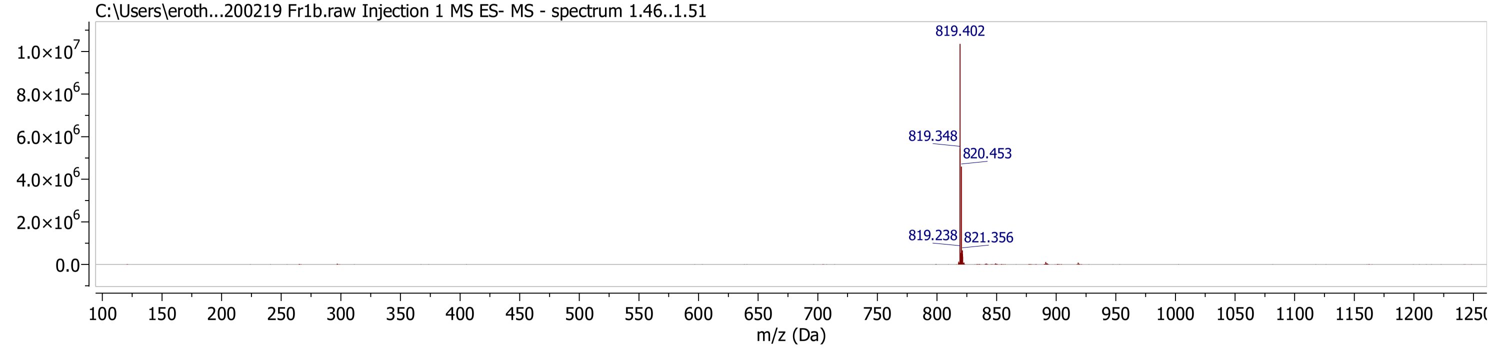
C:\Users\eroth...200219 Fr1b.raw Injection 1 MS ES- MS - spectrum 1.46..1.51
819.402
819.348
820.453
819.238
821.356
m/z (Da)

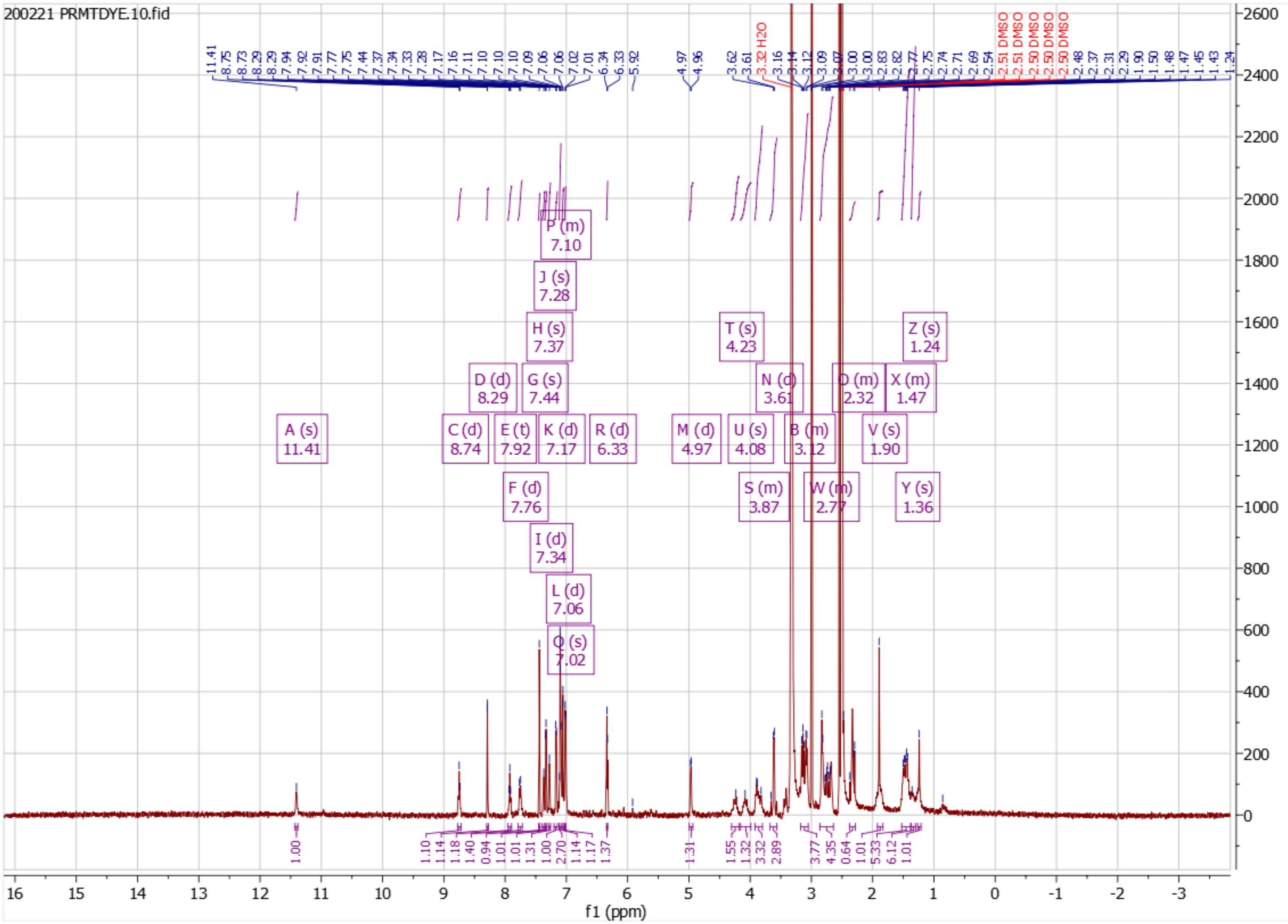

200221 PRMTDYE.10.fid
A (s) 11.41
C (d) 8.74
D (d) 8.29
E (t) 7.92
F (d) 7.76
G (s) 7.44
H (s) 7.37
I (d) 7.34
J (s) 7.28
K (d) 7.17
L (d) 7.06
P (m) 7.10
Q (s) 7.02
R (d) 6.33
M (d) 4.97
T (s) 4.23
U (s) 4.08
S (m) 3.87
N (d) 3.61
B (m) 3.12
W (m) 2.77
O (m) 2.32
V (s) 1.90
X (m) 1.47
Y (s) 1.36
Z (s) 1.24
3.32 H2O
2.51 DMSO
2.51 DMSO
2.50 DMSO
2.50 DMSO
2.50 DMSO
f1 (ppm)
